# Archaeal bundling pili of *Pyrobaculum calidifontis* reveal similarities between archaeal and bacterial biofilms

**DOI:** 10.1101/2022.04.22.489182

**Authors:** Fengbin Wang, Virginija Cvirkaite-Krupovic, Mart Krupovic, Edward H. Egelman

## Abstract

While biofilms formed by bacteria have received great attention due to their importance in pathogenesis, much less research has been focused on the biofilms formed by archaea. It has been known that extracellular filaments in archaea, such as Type IV pili, hami and cannulae, play a part in the formation of archaeal biofilms. We have used cryo-electron microscopy to determine the atomic structure of a previously uncharacterized class of archaeal surface filaments from hyperthermophilic *Pyrobaculum calidifontis*. These filaments, which we call archaeal bundling pili (ABP), assemble into highly ordered bipolar bundles. The bipolar nature of these bundles most likely arises from the association of filaments from at least two different cells. The component protein shows homology, both at the sequence and structural level, to the bacterial protein TasA, a major component of the extracellular matrix in bacterial biofilms, contributing to biofilm stability. We show that ABP forms very stable filaments in a manner similar to the donor-strand exchange of bacterial TasA fibers and chaperone-usher pathway pili where a β-strand from one subunit is incorporated into a β-sheet of the next subunit. Our results reveal mechanistic similarities and evolutionary connection between bacterial and archaeal biofilms, and suggest that there could be many other archaeal surface filaments that are as yet uncharacterized.

**Significance:** Biofilms are communities of microbes where cells attach to each other as well as to surfaces, and bacterial biofilms have been intensively studied due to their importance in many infections. Much less has been known about archaeal biofilms, where archaea are a third domain of life. Using cryo-electron microscopy, we have determined the atomic structure of a surface filament in archaea that forms bi-polar bundles connecting cells. We show that this protein has common ancestry with a protein known to be an important component of bacterial biofilms. This adds to our understanding of the evolutionary relationship between bacteria and archaea and may provide new insights into bacterial biofilms.

## Introduction

In the environment, most microbial species alternate between free-living, planktonic lifestyle, whereby single cells float or swim in liquid medium, and sessile lifestyle, characterized by formation of complex microbial consortia, known as biofilms, in which cells attach to each other as well as to various biotic or abiotic substrates (1-4). It has been even suggested that biofilms represent a default mode of growth for many species, whereas planktonic lifestyle is an artifact of in vitro propagation (5). The cells within a biofilm undergo functional specialization towards a common goal of optimization of the maintenance and viability of the community (1, 6). Thus, different subpopulations within a biofilm are physiologically distinct from each other as well as from planktonic cells, although pinpointing the underlying gene expression pattern remains challenging (7, 8). One important manifestation of these physiological differences is that microbial communities within biofilms display increased resistance to viruses (9) and various antimicrobials, including antibiotics (10, 11). Indeed, studies on biofilms have profound biomedical relevance (12) and hence, most of our understanding of biofilm formation stems from studies on bacterial species.

A common feature of bacterial biofilms is formation of an extracellular polymeric substance, which consists of various polymers, including exopolysaccharides, proteins and extracellular (e)DNA, that are excreted by the cells in the biofilm and encase the bacterial community within a protective matrix (13). In a soil-dwelling, monoderm bacterium *Bacillus subtilis*, one of the most extensively studied biofilm models (14, 15), maintenance of the structural biofilm integrity and the timing of biofilm development depend on a secreted protein TasA (16, 17), which has a globular jelly-roll domain (18) and forms fibrous assemblies (19). TasA is encoded in an operon with *tapA* and *sipW*. A minor matrix component TapA is also a secreted protein and has been described as a TasA assembly and anchoring protein, but its exact function remains unclear (20). The crystal structure of the central TapA domain revealed a β-sandwich fold, most closely related to that of bacterial macroglobulins (21). The third gene in the operon encodes a signal peptidase SipW, which is responsible for the processing of the N-terminal signal peptides in both TasA and TapA (22, 23).

Although biofilms have been most extensively studied in bacteria, archaea are also known to widely practice this lifestyle (24, 25). Indeed, biofilm formation has been documented for multiple metabolically different archaeal species inhabiting highly diverse ecological niches (24, 25). Archaeal biofilms are thought to provide protection against various environmental stresses (26-28), promote horizontal gene exchange (29, 30) and enable syntrophy with other archaea and bacteria (31-34). Molecular studies on archaeal biofilm formation are still in their infancy and have largely focused on genetically tractable hyperthermophilic members of the order Sulfolobales and hyperhalophilic archaea of the class Halobacteria (24). Both Sulfolobales and halophilic archaea deploy different type 4 pili (T4P) during early stages of biofilm formation, whereby cells adhere to various substrates (35, 36). Glycosylation was found to play an important role in this process (37), consistent with the fact that archaeal T4P are typically heavily glycosylated (37-39). The molecular details of archaeal biofilm maturation, involving formation of a complex 3D architecture, is less well understood. In halophilic archaea, exopolysaccharides and eDNA are important components of the extracellular matrix (30, 40), whereas in Sulfolobales biofilms, only small amounts of eDNA were detected (41, 42). Whether fibrous protein assemblies, similar to *B. subtilis* TasA, are involved in the structuring of archaeal biofilms remains unknown.

Here we present a structure of a novel extracellular filament, which we named archaeal bundling pilus (ABP), produced by a hyperthermophilic and neutrophilic archaeon *Pyrobaculum calidifontis* (order Thermoproteales). We show that ABP is homologous to the major biofilm matrix protein TasA of *B. subtilis*. Our structural data explains not only how ABP subunits polymerize through donor-strand exchange, but also how ABP filaments from different cells form antiparallel bundles, which could promote biofilm matrix formation in archaea. These insights might be also relevant to understanding TasA-mediated biofilm formation in bacteria.

## Results

### Bundling filaments at different microscopic scales

*P. calidifontis* is a hyperthermophilic and metabolically versatile archaeon belonging to the order Thermoproteales (phylum Thermoproteota, formerly known as Crenarchaeota) (43). Under scanning electron microscopy (SEM), rod-shaped *P. calidifontis cells* are often seen connected to each other with fibers ranging from 200 to 500 Å in diameter (Fig. 1A). Under negative staining electron microscopy (Fig. 1B,C), these fibers can be resolved to be a mesh formed from thin filaments ∼ 50 Å in diameter, which we name archaeal bundling pili (ABP). The mesh between cells has a single strong axial periodicity of ∼ 150 Å (Fig. 1C, inset) generating the transverse striations that are clearly seen in the micrographs. ABP are seen to directly extend from the cell surface using cryo-electron microscopy (cryo-EM), without any transmembrane secretion complex observed (Fig. 1D). After shearing the filaments off from the cells, ABP remain soluble. Multiple forms of the filaments/bundles are found, ranging from rare single filaments to bundles up to a couple of hundred Ångstroms in diameter (Fig. 1E).

**Fig. 1.**
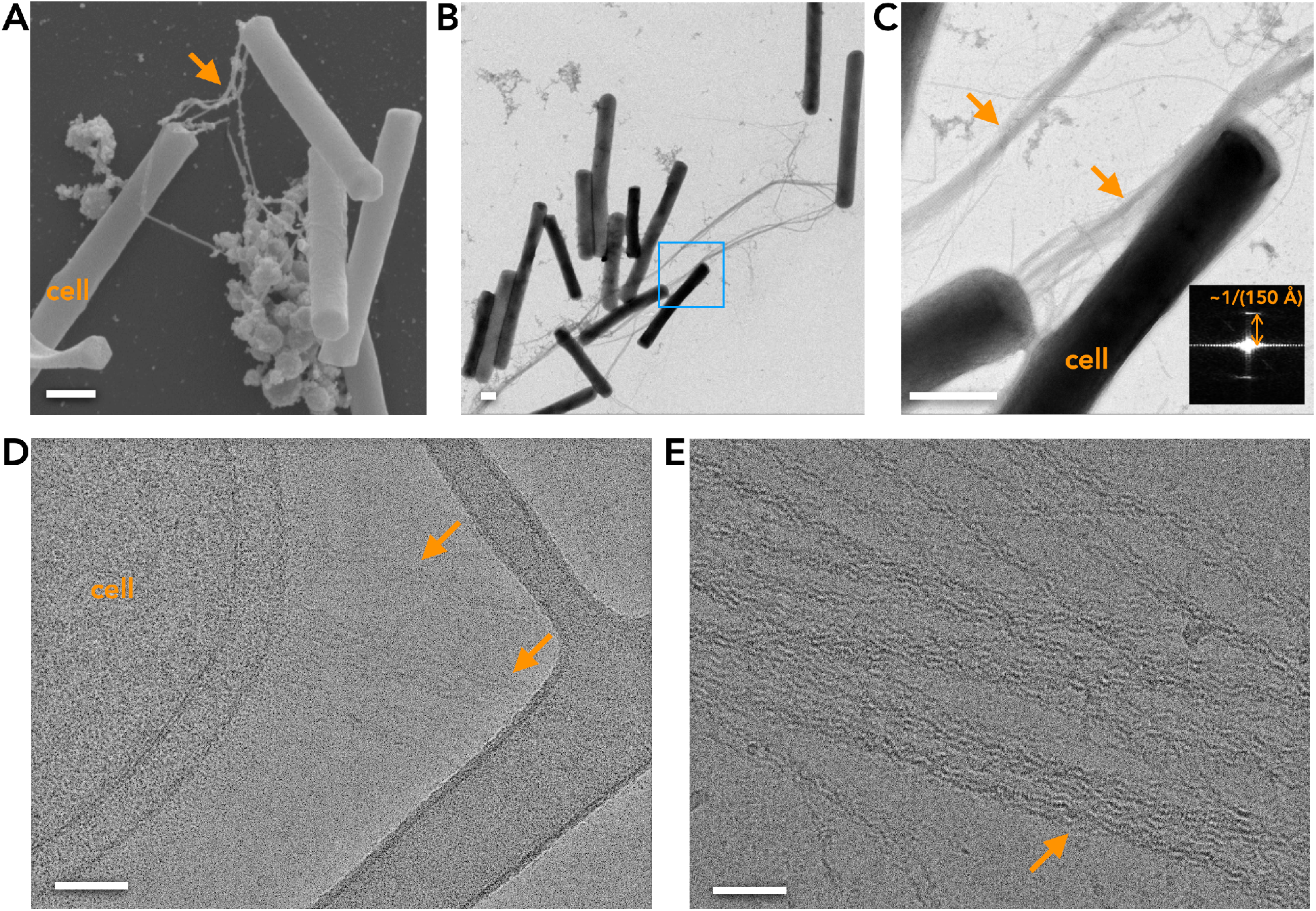
Archaeal bundling pili (ABP) at different scales. (A) Scanning electron microscope (SEM) image of *P. calidifontis* ABP fibers (orange arrow) connecting cells. Scale bar, 500 nm. (B) Negative staining image of ABP fibers connecting *P. calidifontis* cells. Scale bar, 500 nm. (C) Higher magnification image of the boxed area in (B). The orange arrows indicate the dense mesh of ABP fibers seen at this higher magnification. The averaged power spectrum from several regions of this mesh (bottom right) shows a single strong periodicity of ∼ 150 Å. Scale bar, 500 nm. (D) Cryo-EM image of ABP (orange arrow) attached to a *P. calidifontis* cell. Scale bar, 50 nm. (E) Cryo-EM image of ABP bundles (orange arrow) that have been sheared off cells. Scale bar, 50 nm.

### Donor strands inserted in the polymerized ABP

We used cryo-EM to determine the structure of ordered helical bundles of ABP containing six filaments: a central filament surrounded by five outer filaments, and were able to generate an overall 4.0 Å resolution reconstruction as judged by a map:map Fourier shell correlation (FSC; Fig. S1). In these bundles containing helical filaments, the helical symmetry of the central filament (the subunits are related to each other by a left-handed rotation of −77.4° and a translation of 32.8 Å along the helical axis) also relates each of the five outer filaments to each other. This symmetry, a helical pitch of 152.5 Å arising from 4.65 units/turn x 32.8 Å rise per subunit, generates the periodicity of ∼ 150 Å seen in the mesh between cells (Fig. 1C), establishing that the ABP must be the dominant component of this mesh.

We will first focus on the structure of the component filament, and then discuss the structure of the bundle. At this resolution, we were able to manually generate a Cα trace for the protein backbone, with an estimated length of ∼ 175 residues. We were then able to determine the pilin sequence by comparing the cryo-EM map with 148 AlphaFold predictions for proteins encoded by *P. calidifontis* that contained between 175 and 300 residues (44). We used a range that included much larger sequences in case there was a cleaved signal sequence that would not be found in the processed protein. There was only one good match to the map from these 148 predictions, Pcal_0910 (WP_011849593), a 204 aa-long protein. The first residue of Pcal_0910 in the ABP is T25, suggesting that the pre-protein is N-terminally processed prior to the ABP assembly. Indeed, SignalP analysis (45) has shown that Pcal_0910 carries a cleavable signal peptide, with the predicted signal peptidase I cleavage site, 22-AMA↓TL-26, coinciding perfectly with the determined structure. The structure of the ABP subunit is β-strand rich and forms a variant of the jelly-roll fold (Fig. 2A). Similar to chaperone-usher (46-48) and type V pili (49), ABP also has a donor strand inserted into a groove in the β-sheet of an adjacent subunit (Fig. 2). The donor-strand corresponds to the B strand of the canonical jelly-roll fold, creating the BIDG β-sheet and extending this sheet by connecting it with a β-hairpin (see below), which is an additional structural element compared to the typical jelly-roll folds.

**Fig. 2.**
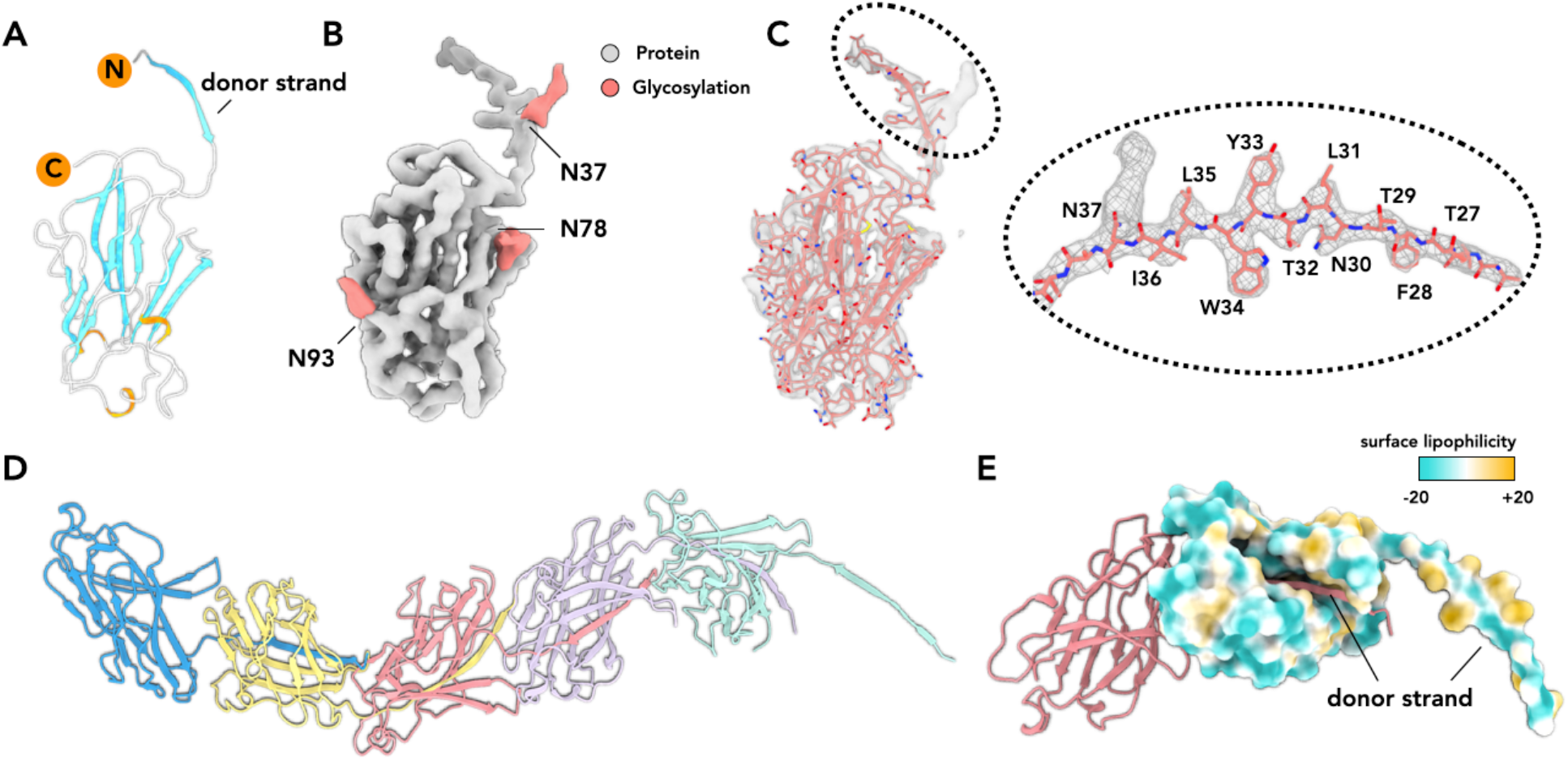
The archael bundling pilin inserts a donor strand into the adjacent pilin. (A) A ribbon representation of a single *P. calidifontis* bundling pilin, with N- and C-terminii labeled. The β-strands are in cyan, and three very short α-helices are in orange. (B) The density accounted for by the pilin atomic model is in gray, and the extra density, due to post-translational modifications, is in red. Three putative glycosylation sites are N37, N78 and N93. (C) The cryo-EM map of a single pilin subunit. The close-up view of the donor strand is shown on the right. (D) The filament model of ABP. Each subunit is in a different color. (E) Two adjacent subunits in the ABP. The left one is shown in a ribbon representation, and the right one is shown in an atomic surface view and colored by lipophilicity.

Extra densities were observed on three asparagines, including one on the donor strand (Fig. 2B-C), all of which are part of an Asn-X-Ser/Thr motif (where X is any amino acid) for N-linked glycosylation (50). The interface between two adjacent subunits (Fig. 2D) is 2,100 Å^2^, as calculated by PDB-PISA, which is quite large compared to the size of the pilin. The surface of the pilin is predominantly hydrophilic, while part of the donor strand and the groove in which it binds is more hydrophobic (Fig. 2E).

### Filaments within the ordered ABP bundles have opposing polarities

We asked how filaments within the ABP bundle, with a total diameter of around 170 Å (Fig. 3A), interact with each other. Within any cross-section perpendicular to the helical axis, the central filament only makes contact with one outer filament, consistent with the helical twist of the bundle with 4.65 subunits per turn (Fig. 3B), which is the same twist as in the component filaments. The outer filaments make no contacts with each other, so the bundle is held together exclusively by contacts with the central filament. A local resolution estimation (Fig. 3C) shows that the central filament has a higher resolution than determined by the overall FSC, at better than 4 Å. In contrast, for the outer filaments, only the subunits making contact with the central filament have a resolution close to 4 Å, and the remainder of the outer filaments are at a lower resolution, presumably due to flexibility.

**Fig. 3.**
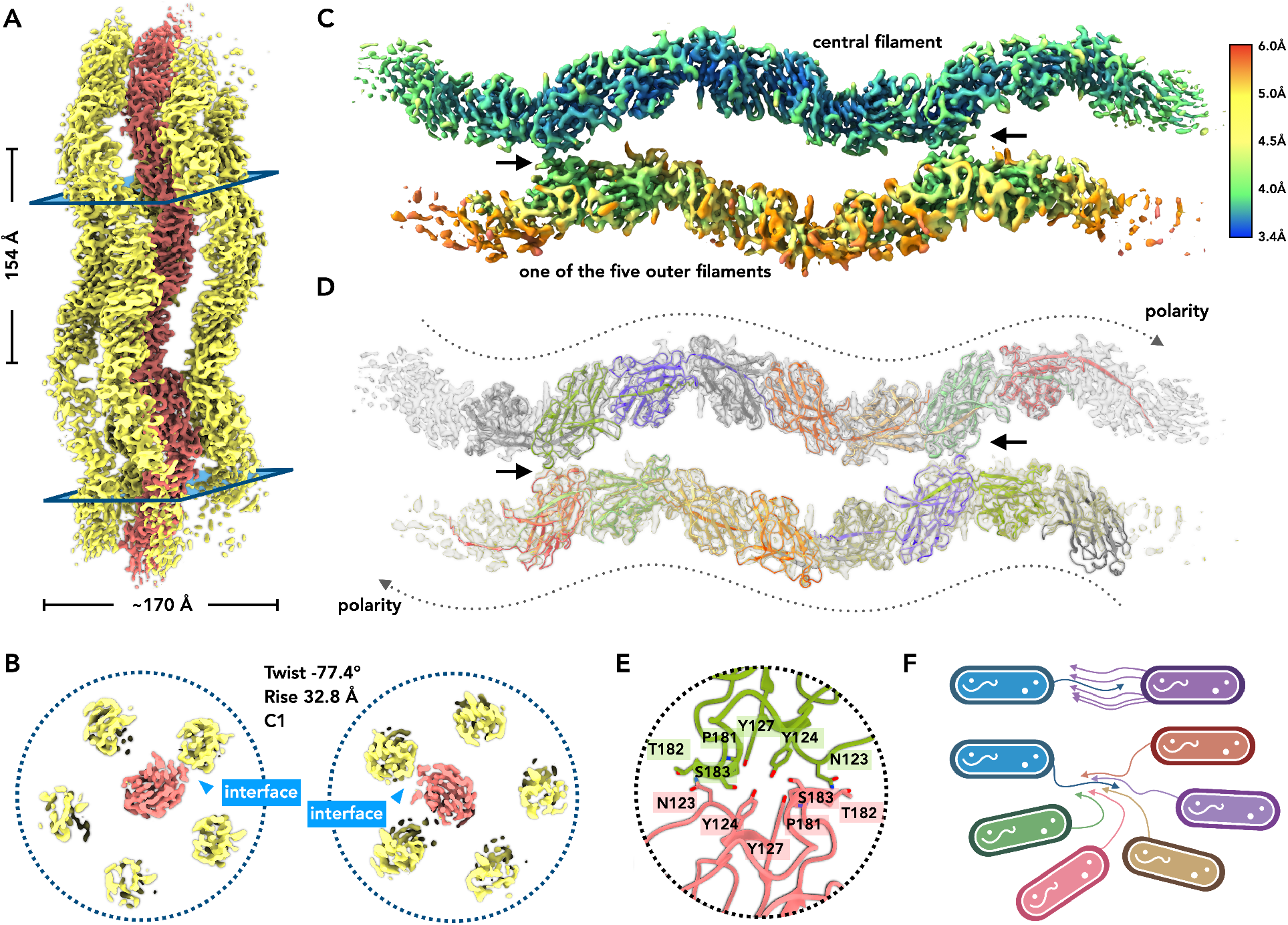
Filaments in the ABP bundle have opposing polarities, elucidating biofilm formation. (A) Cryo-EM map of ABP. A global helical symmetry exists in the six-filament bundle, with the central filament in red and the other five filaments in yellow. The diameter of the six-filament bundle is ∼ 170 Å, and the pitch of this bundle is 154 Å. (B) The filament-filament interface between the central filament and the outer filaments, shown at the level of the two planes indicated in (A). (C) The local resolution estimation of the central filament and one of the five outer filaments. The filament-filament interfaces are indicated by black arrows. (D) The same view as (C), with atomic models of these two filaments shown in the map. The map shows that these two filaments have opposite polarities. (E) A close-up view of the filament-filament interface, with the residues involved in the interface labeled. The central filament is in green, and the outer filament is in pink. (F) A schematic model for the role of the bundling pili in biofilm formation. Such a bundle can be formed between as few as two cells (top), but it can also connect, in principle, up to six cells within one bundle (bottom).

What is most striking in the bundle is that the outer filaments have the opposite polarity compared to the central filament, suggesting that they probably come from different cells (Fig. 3D), consistent with the electron microscopy observations showing that ABP bundles are anchored within the envelopes of interconnected cells (Fig. 1B,C). Each interface between an outer filament and the central filament is mediated mainly by hydrogen bonds, with a small buried interfacial area of 400 Å^2^ (Fig. 3E). Due to the bipolar nature of the bundle, this interface has a local pseudo-C2 symmetry (Fig. 3E) and repeats every five subunits. With this bundling mechanism, one *P. calidifontis* cell can form a robust bridge with another cell and have the potential to connect up to five different cells simultaneously (Fig. 3F).

### ABP is a homolog of bacterial TasA

We next assessed the evolutionary provenance of Pcal_0910 by performing sensitive profile-profile comparisons using HHpred (51) against the protein family database (Pfam). The search yielded a highly significant (97% probability) hit to a profile hidden Markov model of Peptidase_M73 protein family (PF12389), which includes camelysin, a secreted metallo-endopeptidase (52), and TasA, a key component of bacterial biofilms (18, 53-58). Indeed, it has been suggested that TasA has evolved from an inactivated camelysin-like protease (18). The match encompassed the N-terminal signal sequence and the C-terminal β-strand rich ectodomain of both proteins (Fig. S2). While the Dali server (59) failed to find significant structural similarity between ABP and TasA, possibly, at least in part, due to donor-strand insertion in the APB but not in the crystalized monomeric TasA (PDB id: 5OF1) (18), manual comparison of the two proteins revealed a high degree of similarity in the jelly-roll domain of both (Fig. 4). Furthermore, a very recent cryo-EM study reveals globular TasA subunits arranged into a filament with a donor strand from one subunit inserted into a sheet in the next subunit (60), very similar to what we report herein for ABP. Collectively, these sequence and structural comparisons establish an evolutionary relationship between ABP and TasA.

**Fig. 4.**
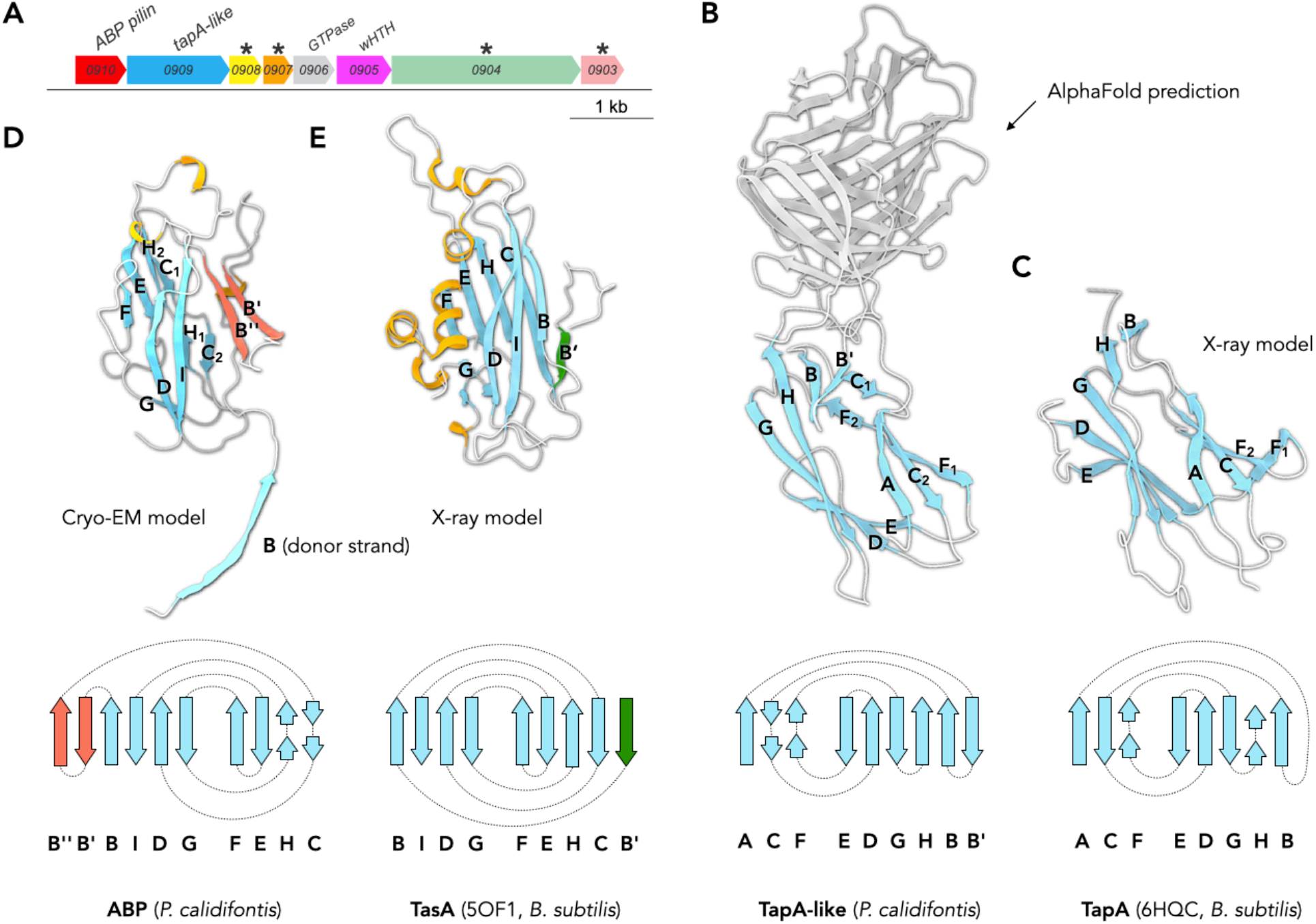
ABP is a homolog of bacterial TasA. (A) Genes in the conserved *Pyrobaculum* operon, including TasA-like and TapA-like genes. Asterisks denote ORFs with predicted transmembrane domains. (B) The AlphaFold prediction of *P. calidifontis* TapA-like protein. The domain homologous to bacterial TapA is highlighted and the schematic representation of the fold is shown below. (C) The TapA protein of *B. subtilis* determined by X-ray crystallography (PDB 6HQC), and its corresponding schematic representation. (D) Cryo-EM model of *P. calidifontis* ABP has a jelly-roll fold, with an additional β-hairpin (red). (E) The X-ray model of *B. subtilis* TasA (PDB 5OF1), which also has a jelly-roll fold.

We further explored the genomic neighborhood similarities of TasA and APB. As mentioned above, in *B. subtilis*, TasA is encoded within an operon together with a putative anchoring protein TapA and a signal peptidase SipW (15). ABP is conserved in hyperthermophilic archaea of the orders Thermoproteales and Thermofilales. In Thermoproteales, the pilin is encoded within a potential operon including eight genes (Fig. S3). In *P. calidifontis*, immediately downstream of Pcal_0910 is a gene encoding a 410 aa-long protein (Pcal_0909) with a predicted signal peptide. The protein does not have recognizable homologs outside of Thermoproteales and Thermofilales. However, AlphaFold modeling of this protein revealed three β-strand rich domains, with the first, signal peptide-proximal β-sandwich domain showing remarkable structural similarity to the ectodomain of the *B. subtilis* protein TapA (Fig. 4A-C), the putative anchoring component for the extracellular TasA filaments (21, 53). Both Pcal_0909 and TapA homologues have a distinct β-sandwich fold, different from the jelly-roll domain of TasA and ABP (Fig. 4D-E). Analysis of the six other proteins encoded within the *P. calidifontis* ABP operon yielded only generic functional predictions. Namely, Pcal_0906 is a predicted GTPase (not conserved in the operons of other Thermoproteales species; Fig. S3) and Pcal_0905 is a winged helix-turn-helix domain containing transcription regulator. Notably, Pcal_0908 (WP_193322921), Pcal_0907 (WP_011849590) and Pcal_0903 (WP_193322920) each have four predicted transmembrane domains, with an N-terminal signal sequence, whereas Pcal_0904 (WP_011849587) has a single N-terminal transmembrane domain (Fig. S4) and a large β-strand rich ectodomain. These proteins might also participate in biofilm formation and/or anchoring of the ABP. In members of the Thermofilales, ABP-encoding loci differ considerably from those of Thermoproteales, with only ABP pilin and TapA-like protein genes being shared between the two types of loci.

### Archaeal TasA homologs beyond Thermoproteales and Thermofilales

The relationship between the ABP and TasA is recognizable using sensitive profile-profile and structural comparisons. We inquired whether archaea encode TasA homologs which would be more readily recognizable at the sequence level. To this end, we assessed the distribution of the Pfam PF12389 family in bacterial and archaeal genomes available in the Genome Taxonomy Database (GTDB) using AnnoTree (61, 62) (Supplementary Data 1). In bacteria, TasA homologs are widespread but not universally conserved. They are readily identifiable in different Firmicutes lineages as well as in several other large phyla, including Actinobacteriota, Chloroflexota, Fusobacteriota and Patescibacteria, whereas diderm bacteria of the phylum Proteobacteria nearly completely lack TasA homologs (Fig. 5A). Notably, we also found TasA homologs to be relatively widespread in archaea, especially, in several lineages previously constituting the phylum Euryarchaeota (recently redistributed into phyla Halobacteriota, Methanobacteriota and Thermoplasmatota) (63). These include members of the orders Halobacteriales, Methanosarcinales, Archaeoglobales, Methanonatronarchaeales, Methanomassiliicoccales and Thermococcales as well as several unclassified lineages (Fig. 5B). In Thermoproteota (former Crenarchaeota), only Caldarchaeales and an unclassified lineage B26-1 encode TasA homologs. Archaeal proteins showing closer similarity to bacterial TasA homologs than ABP display a patchy distribution suggestive of extensive horizontal gene transfer. Although the function of these archaeal protein remains to be investigated experimentally, it is likely that they are also implicated in biofilm formation, further suggesting that the mechanism of biofilm structuring in bacteria and, at least, some archaea depends on homologous proteins.

**Fig. 5.**
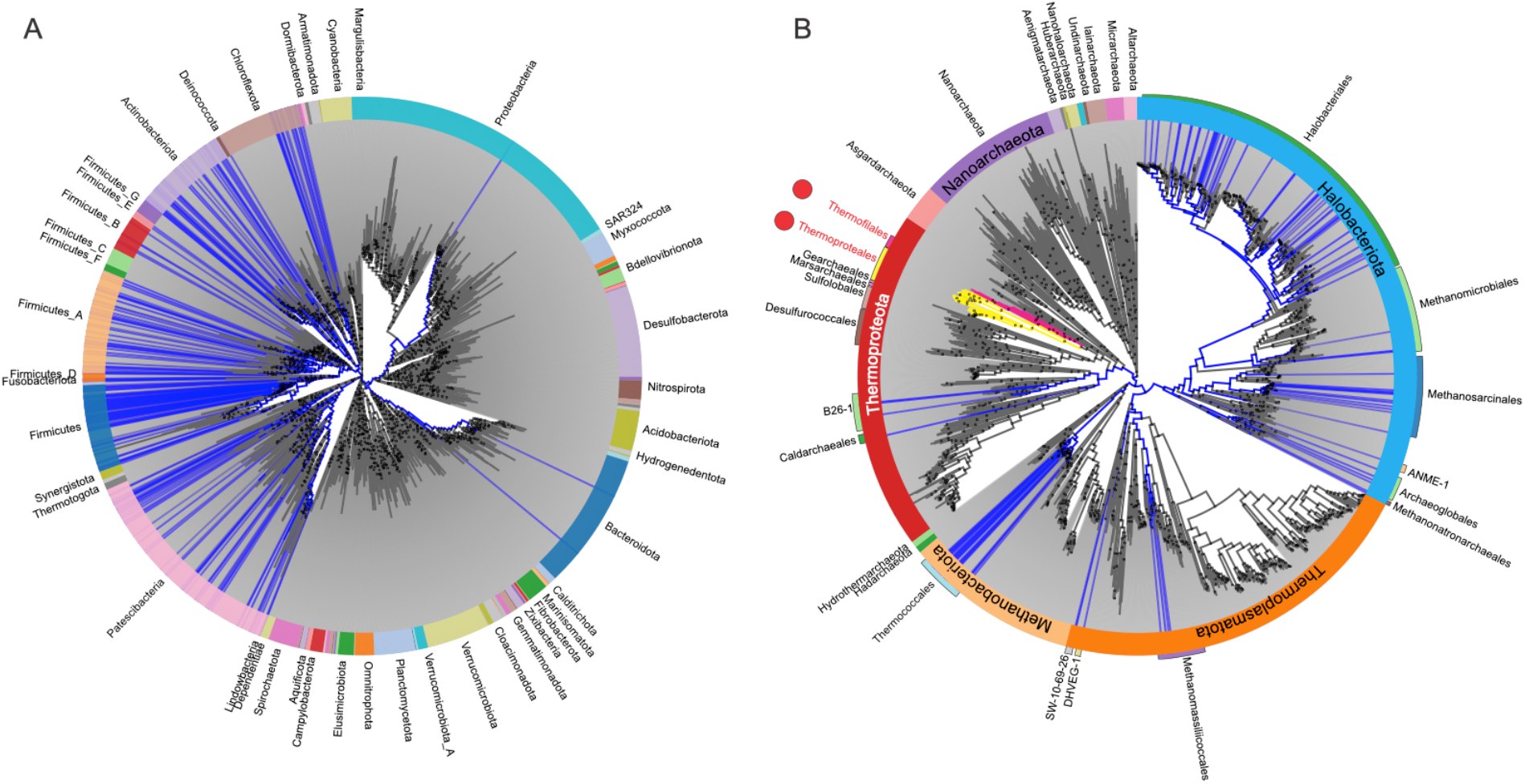
Phylogenetic distribution of TasA-like proteins in bacterial (A) and archaeal (B) genomes. Branches corresponding to bacterial or archaeal species which encode TasA-like proteins matching the Pfam profile PF12389 (E-value cutoff of 0.005) are shown in blue, whereas those lacking it are in black. Branches of Thermoproteales and Thermofilales which encode more divergent TasA homologs are shown in yellow and magenta, respectively. The trees are scaled to show family and species level diversities for bacteria and archaea, respectively. The outer circle indicates the phyla, which are labeled. Relevant archaeal orders are also shown as bars. The figure was prepared using AnnoTree (http://annotree.uwaterloo.ca). Sequences of bacterial and archaeal TasA-like proteins are provided in Supplementary Data 1.

## Discussion

There is an extensive literature detailing how pili in the chaperone-usher pathway use a mechanism of strand exchange to form filaments with interlocking β-sheets (64). For these pilin subunits, a chaperone provides a strand to complete a β-sheet in the pilin prior to assembly. The usher in the outer membrane has the catalytic activity to exchange the strand from the chaperone with a donor strand from another subunit, forming the filament. Since the structure of ABP has a formal resemblance to that of chaperone-usher pili, with a donor strand from one subunit inserted into a β-sheet in the next subunit, we must ask whether a chaperone and usher are also involved. Analysis of the *P. calidifontis* genome does not detect any obvious candidates for either a chaperone or a catalytic usher. Indeed, comparison of the ABP pilin encoding operons in Thermoproteales and Thermofilales showed that only two genes, namely, the pilin gene itself and a gene encoding a TapA-like protein, are conserved between the two groups of archaea (Fig. S3).

An AlphaFold prediction for the structure of the ABP monomer (Fig. S5) has the donor strand folding back into the subunit, to self-complement. So it is possible that such self-complementation takes place, eliminating the need for a chaperone to maintain the fold of the subunit prior to assembly. But it is hard to imagine how such a self-complemented subunit could remove the donor strand and insert it into another subunit, displacing the strand there, in the absence of some complicated catalytic pathway. Since the donor strand is rather hydrophobic (Fig. 2E), it is possible that the strand partially folds back on the subunit, but is not inserted into the sheet as in the AlphaFold model, rather than projecting out as in the diagrams of Fig. 2A-C. If that were the case, glycosylation at N37 on the donor strand and N78 near the gate of the groove might shield the hydrophobic residues and keep the donor strand soluble without self-inserting. The latter case would lead to a much smaller energy barrier when the pilin assembles into a filament, as the strand would not need to be removed from the sheet in the donor subunit to insert into the next subunit. Notably, the donor-strand (strand B) not only reconstitutes the BIDG β-sheet of the canonical jelly-roll fold but also extends this β-sheet to include a β-hairpin formed by B’ and B’’ β-strands (Fig. 4D), further stabilizing the ABP filament. The archaeal protein CanA (65) might also provide some insights into assembly of ABP. It has been shown using a purely *in vitro* system that CanA self-assembles into filaments (PDB: 7UII) in the absence of any chaperone or catalyst. In the filament, a donor strand from one subunit is inserted into a β-sheet of another subunit. Thus, CanA might serve as a model for a self-assembly process that does not require a chaperone or catalytic usher as is required in the bacterial chaperone-usher pathway.

Our results have uncovered an unexpected evolutionary relationship between the loci encoding ABP and TasA, a major matrix protein orchestrating biofilm formation in bacteria. We show that not only are the major pilins homologous at the sequence and structural level, but the minor component of the bacterial biofilm matrix, protein TapA, also has a counterpart in archaea. Whereas in bacteria the signal peptides of both TasA and TapA are processed by a dedicated signal peptidase, SipW, encoded in the *tapA*-*sipW*-*tasA* operon, proteolytic processing of the archaeal counterparts appears to be performed by the generic signal peptidase as no dedicated gene was identified in the vicinity of the ABP operon. The ABP and TapA-like proteins conserved in archaeal orders Thermoproteales and Thermofilales display considerable divergence compared to their bacterial homologs, with the relationship being detectable only using some of the most sensitive profile-profile or structural comparisons. However, we found that proteins more closely related to the bacterial TasA homologs display rather broad, even if patchy, distribution in archaea, suggesting that TasA-mediated biofilm formation might be more widespread across both prokaryotic domains of life than currently appreciated. Future functional and structural studies on these TasA-like proteins (Supplementary Data 1) are bound to uncover further parallels between bacterial and archaeal biofilms. It has been previously considered that T4P are the only cell surface filaments shared between bacteria and archaea (24, 38, 39, 66, 67). Our current results extend the similarities between bacterial and archaeal surface structures to TasA/ABP. It is almost certain that further research on archaeal cell envelopes will uncover additional surface filaments, both archaea-specific and those shared with cellular organisms from the other domains of life.

Our structure of the *P. caldidifontis* ABP bundle provides unique insights into how ABP filaments emanating from different cells can promote cellular aggregation and biofilm formation. In particular, bipolar organization of the bundle can entangle up to six individual cells (Fig. 3F). Notably, negative staining electron microscopy observation of ABP bundles (Fig. 1B) suggests that the bundles may include more than six filaments and hence more cells can be envisioned to participate in such interactions. Bundles composed of multiple TasA fibers have been observed in *B. subtilis* biofilms. However, high-resolution structural data for such bundles is not available at the moment, with only doublets of TasA filaments having been investigated using cryo-electron tomography-based modeling and molecular dynamics simulation (60). The bipolar ABP structure presented in this work might be relevant for understanding the formation of larger bacterial TasA bundles. Collectively, our results uncover previously unsuspected mechanistic similarities and evolutionary connections underlying the formation of bacterial and archaeal biofilms.

## Materials and Methods

### Cell growth and pili purification

*Pyrobaculum calidifontis* DSM 21063 cells (43) were purchased from the DSMZ culture collection and grown in 1090 medium (0.1% yeast extract, 1.0% tryptone, 0.3% sodium thiosulfate, pH7) at 90 °C without agitation. 30 ml pre-culture was started from a 200 µl cryo-stock, grown for two days and then diluted into 200 ml of fresh medium. When OD_600_ reached ∼0.2, the cells were collected by centrifugation (Sorval SLA1500 rotor, 7000 rpm, 10 min, 20 °C). The resultant pellet was resuspended in 4 ml of PBS buffer and the cell suspension was vortexed for 15 min to shear off the extracellular filaments. The cells were removed by centrifugation (Eppendorf F-35-6-30 rotor, 7830 rpm, 20 min, 20 °C). The supernatant was collected and the filaments were pelleted by ultracentrifugation (Beckman SW60Ti rotor, 38000 rpm, 2h, 15 °C). After the run, the supernatant was removed and the pellet was resuspended in 200 µl of PBS.

### Scanning electron microscopy

The *P. calidifontis* cells were grown in 1090 medium at 90 °C without shaking until OD_600_ reached ∼0.2. Then, 18 ml of cell culture was fixed by adding 2 ml of 25 % glutaraldehyde solution, incubated at room temperature for 2 h and kept overnight at 4 °C. The fixed culture was then centrifuged (Eppendorf F-35-6-30 rotor, 7000 rpm, 10 min, 20 °C) and the cell pellet was resuspended in 2 ml of 0.1 M HEPES, 2.5 % glutaraldehyde, pH 7.2 buffer. One ml of cell suspension was placed on poly-L-lysine-coated coverslip and incubated overnight at 4 °C. Then the cells were washed three times with 0.1 M HEPES buffer pH 7.2, post-fixed for 1 h 30 in 1% osmium tetroxide in 0.1 M HEPES pH 7.2 and rinsed with distilled water. Samples were then dehydrated by gradually washing in ethanol solutions (25, 50, 75, 95 and 100%), followed by critical point drying with CO_2_. Dried specimens were sputtered with 20 nm gold palladium using a GATAN Ion Beam Coater and imaged with a JEOL JSM 6700F field emission scanning electron microscope.

### Transmission electron microscopy

For negative staining transmission electron microscopy, 10 µl of cell suspension were adsorbed onto glow-discharged copper grids with carbon-coated Formvar film and negatively stained with 2.0% (w/v) uranyl acetate. The samples were observed under FEI Tecnai Spirit BioTwin 120 microscope operated at 120 kV.

### Cryo-electron microscopy and image processing

The *P. calidifontis* sample was applied to glow-discharged lacey carbon grids and vitrified using a Leica EM GP plunge freezer (Leica). Grids were imaged on a Titan Krios (300 keV, Thermo Fisher) with a K3 camera (Gatan). 2,027 micrographs were collected under electron counting mode at 1.08 Å per pixel, using a defocus range of 1–2 μm with ∼50 electrons/Å^2^ distributed into 40 fractions. Motion correction and CTF estimation were done in cryoSPARC (68-70). Initial 2D references were generated from ∼200 manually picked particles and these were used for autopicking 380,371 segments. From these segments, 2D class averages of bundles with high-resolution details were found, and the segments classified by these averages were used for a helical reconstruction (Fig. S6). The two possible helical symmetries were calculated from an averaged power spectrum generated from the raw particles and one was found to be correct (71, 72). Typically, the hand of a cryo-EM protein reconstruction is determined by inspecting the hand of α-helices in the map (73). However, this map only has two tiny one-turn helices that were not obvious without an atomic model. So, the map hand was initially determined by comparing with the AlphaFold prediction, and later validated by the hand of two tiny α-helices. The resolution of the reconstruction was estimated by both Map:Map FSC, Model:Map FSC, and local resolution estimation in cryoSPARC. The statistics are listed in supplemental Table 2.

### Protein identification and model building

The density corresponding to a single bundling pilin was segmented from the experimental cryo-EM density using Chimera (74). Manual Cα tracing estimated the subunit to have ∼175 amino acids. A total of 148 protein structures with lengths between 175 and 300 in *P. calidifontis* were predicted by AlphaFold (44). All predictions were then compared with the Cα trace for the best match, and the protein WP_011849593 was identified as the correct pilin. This protein identification approach has been developed into an online tool, DeepTracer-ID (paper submitted). The full-length sequence was threaded into the map using DeepTracer (75), manually adjusted in Coot (76), and real-space refined in PHENIX (77). MolProbity was used to evaluate the quality of the filament model (78). The refinement statistics are shown in Table 1.

**Table 1.** Cryo-EM of *P. calidifontis* bundling pili.

| Parameter | Bundling pili |
| --- | --- |
| **Data collection and processing** |  |
| Voltage (kV) | 300 |
| Electron exposure (e<sup>-</sup> Å<sup>-2</sup>) | 50 |
| Pixel size (Å) | 1.08 |
| Particle images (n) | 204,565 |
| Shift (pixel) | 40 |
| **Helical symmetry** |  |
| Point group | C1 |
| Helical rise (Å) | 32.79 |
| Helical twist (°) | -77.40 |
| **Map resolution (Å)** |  |
| Map:map FSC (0.143) | 4.0 |
| Model:map FSC (0.38) | 4.1 |
| d<sub>99</sub> | 4.4 |
| **Refinement and Model validation** |  |
| Ramachandran Favored (%) | 91.2 |
| Ramachandran Outliers (%) | 0.2 |
| RSCC | 0.79 |
| Clashscore | 18 |
| Bonds RMSD, length (Å) | 0.028 |
| Bonds RMSD, angles (°) | 0.731 |
| **Deposition ID** |  |
| PDB (model) | 7UEG |
| EMDB (map) | EMD-26474 |

### Sequence analyses

Profile-profile comparisons were performed using HHpred against the Pfam database (51). Signal peptides and transmembrane domains were predicted using SignalP v5 (45) and TMHMM (79), respectively. Genomic neighborhoods were analyzed using the enzyme function initiative– genome neighbourhood tool (EFI-GNT)(80) and SyntTax (81). The distribution of TasA like proteins (Pfam PF12389) in the bacterial and archaeal genomes available in the Genome Taxonomy Database (GTDB) was assessed using AnnoTree (61, 62).

## Data Availability

The cryo-EM atomic model and map have been deposited in the PDB (ID code 7UEG) and Electron Microscopy Data Base (ID EMD-26474), respectively.

## Acknowledgements

The cryo-EM imaging was done at the Molecular Electron Microscopy Core Facility at the University of Virginia, which is supported by the School of Medicine. This work was supported by NIH Grant GM122510 (E.H.E.) and K99GM138756 (F.W.). The work in the M.K. laboratory was supported by grants from l’Agence Nationale de la Recherche (ANR-20-CE20-009-02 and ANR-21-CE11-0001-01) and Ville de Paris (Emergence(s) project MEMREMA). We are grateful to Ultrastructural BioImaging Core Facility of Institut Pasteur for the access to electron microscopes and Christine Schmitt for help with sample preparation for SEM.

