## Supplemental appendix for "Archaeal bundling pili of *Pyrobaculum calidifontis* reveal similarities between archaeal and bacterial biofilms"

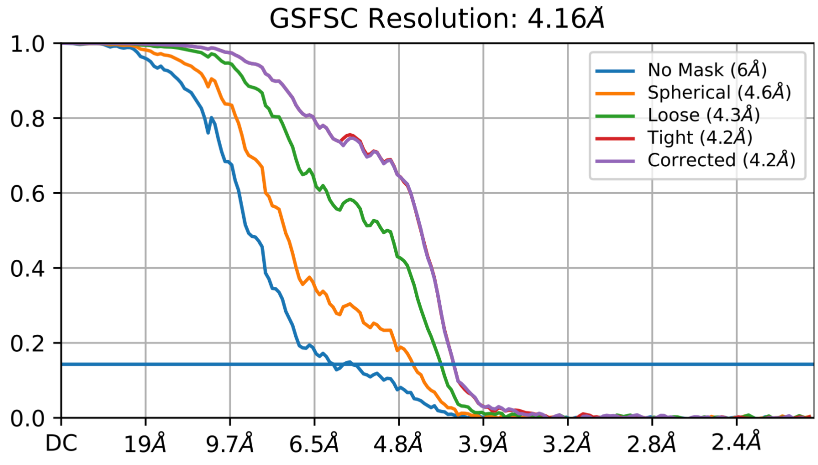


**Fig. S1**. Fourier Shell Correlation (FSC) calculation

The map:map "gold standard" FSC using the 0.143 criterion estimates the final reconstruction to have a global resolution of 4.2 Å.

```
PF12389.11 ; Peptidase_M73 ; Camelysin metallo-endopeptidase/TasA
Probability: 97.21%,  E-value: 0.11,  Score: 43.17,  Aligned cols: 162,  Identities: 13%,  Similarity: 0.054,  Template Neff:
10.3

Q ss_pred                   CccccccHHHHHHHHHHHHHHHHHHHHHhhceeeeE--EEeeCCCeEEecCCCCCccccccCCcEEEEEEEecCCCCcc
Q WP_011849593.1    1    MARKKNYRPLIALAALAVAALAMATLTFTNLTYWL--INATLPPAMKYPGTDTTITRSDSSGYNRYVYVSYYYDPSTGYN   78 (204)
Q Consensus         1    ~~~~k~~~~l~ll~~~l~~a~a~AaV~~~n~~~~~~--V~~~~pPV~~~~G~d~~~~g~dl~~~g~~i~v~~~~~~~~G~n   78 (204)
                         |||||++..++++++++++....+.-++.......  +++....+.+......
T Consensus         1    ~~~k~~~~~~~~~~~~~~~~~~t~A~~s~~~~~~~~~~~~g~l~i~~~~~~~~~---------------------------   53 (193)
T PF12389.11        1    TLKKKLGMGIASAVLGAALVGGGTFAFFSDKEVSNNTFATGTLDLALNPSTVV---------------------------   53 (193)
T ss_pred                   CchHHHHHHHHHHHHHHHHHHHHHHHHHHcccccceeeEEEEeEEEEEcCCCceee---------------------------

Q ss_pred                   eEEEEEEEe-CCCcEEEcceEEEecCCCCCcE-EEEEEEEeccCCC-----CccccceEEEEEEEECCCCCCe-EEEEcCc
Q WP_011849593.1    79   VTRISIVGF-TGDPTNYTNVLQLCNKYYSGTL-YAKLVAVGTVGTT-----NYESYIKDFRVYFVNPTTTPN-YVQFQGT   150 (204)
Q Consensus         79   ~T~isi~~~-tg~~~~Y~~vl~I~N~d~s~~~~-v~L~~~~~~~~~~~~~-----g~~~yv~~i~l~i~~~~g~~~~-v~~~~t   150 (204)
                         .+.-.  ||+....  .+.|.|..  +...  +++|+........  +  ...+.+.+.+........  .+....-
T Consensus         54   ----~~~~~~PG~~~~~~~~~~v~N~g-~~~~~~~~~~l~~~~~~~~~~~~~~~~~--~l~~~l~~~~~~~~~~~~~~~~~~~~   124 (193)
T PF12389.11        54   ----NVSNLKPGDTVEK--EFKLENKG-TLDIKKVLLKTDYNVEDVKKDNKD--DFGKHIKVTFLKNVDKHETIVKETAL   124 (193)
T ss_pred                   ----eccccCCCCEEEE--EEEEEeCC-CCCceEEEEEEEeeecCCCCCCCcc--hhhhheEEEEEeecCCCcceEEeeee

Q ss_pred                   cccccccce-----------------EEECCCCEEEEEEEEEECCCC---CCccccCCCeEEEEEEEEEEEecCC
Q WP_011849593.1    151  SVTQSATGS-----------------VSIGPGQCATVGAYVLVDPSL---PTSARDGKTVIATYQVNVVFSTSP   204 (204)
Q Consensus         151  ~~~~s~t~~------------------~tI~aG~~~~idvev~i~~~a---~~~~~~~~g~~~~~~~~~qvdV~~~~~~~   204 (204)
                         .  .-..  ..|++|+...+.+.+.++..+  ...  .+....+.++|+.+..+.+
T Consensus         125  ----~~~~~~~~~~~~~~~~~~~~~~~~~l~~g~~~~~~~~~~~~~~~~~~~~~~n~~~~qg~~~~~~~~~~~~a~Q~~~~   192 (193)
T PF12389.11        125  D----KLKGDTLTAVNNDLAAWFWDEKGISAGKSDKFKVKFEFVDNKKDQNEF--QGDKLQLTWTFDAQQGDGE   192 (193)
T ss_pred                   e----hhhcCchhhhccccchhhcccCCCCCCCCCCeEEEEEEEcCCCCCCccee--ccCEEEEEEEEEEEEecCCC
```

**Fig. S2.** Results of the HHpred analysis queried with the Pcal_0910 (WP_011849593) sequence. H(h), α-helix (red); E(e), β-strand (blue); C(c), coil (grey).

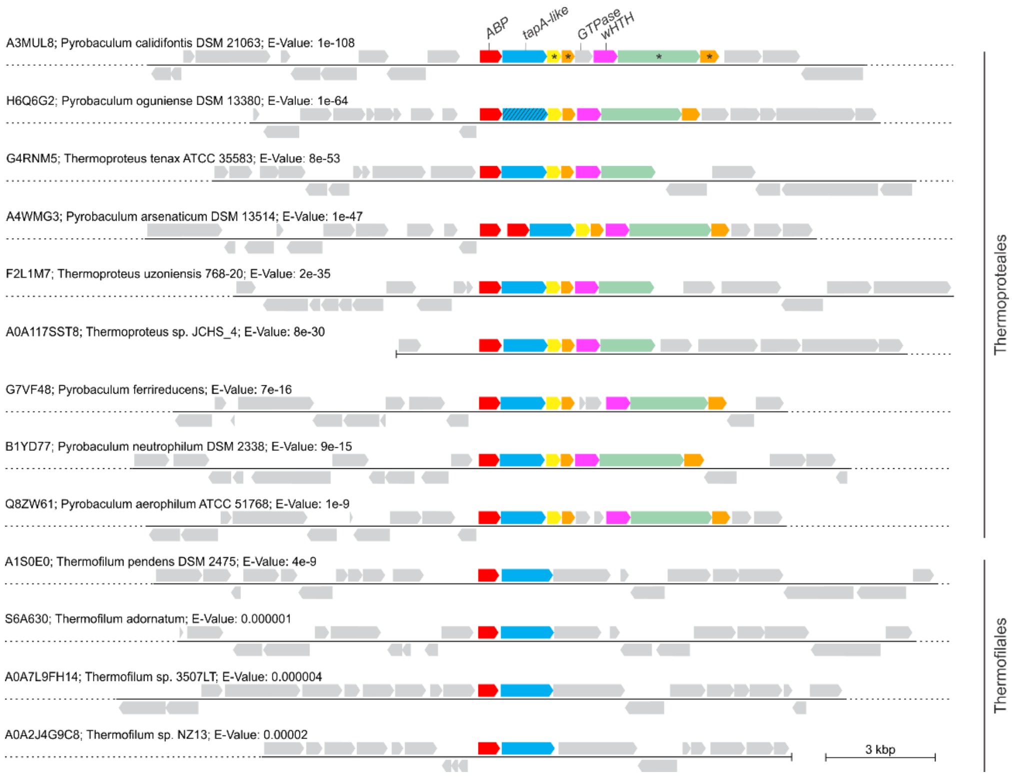


**Fig. S3.** Genomic loci encoding ABP pilins in the genomes of Thermoproteales and Thermofilales. Conserved genes in the predicted ABP operon of Thermoproteales are color coded, with homologous genes in the same color. Genes surrounding the ABP locus are shown in grey. In *P. oguniense*, the TapA-like gene is inactivated and is indicated with hatched lines.The corresponding loci in Thermofilales include only two genes, for ABP pilin (red) and TapA-like protein (blue), homologous to those in Thermoproteales. Genomic loci are aligned using the ABP pilin genes (red) and indicated with the corresponding UniProt accession numbers, followed by the organism name and E-value of the blastp hit to the reference ABP pilin of *Pyrobaculum calidifontis*. Asterisks denote ORFs with predicted transmembrane domains. Abbreviations: ABP, archaeal bundling pilus; GTPase, guanosine triphosphate hydrolase; wHTH, winged helix-turn-helix domain-containing protein. Genomic neighbourhoods were analysed using the enzyme function initiative–genome neighbourhood tool (EFI-GNT).

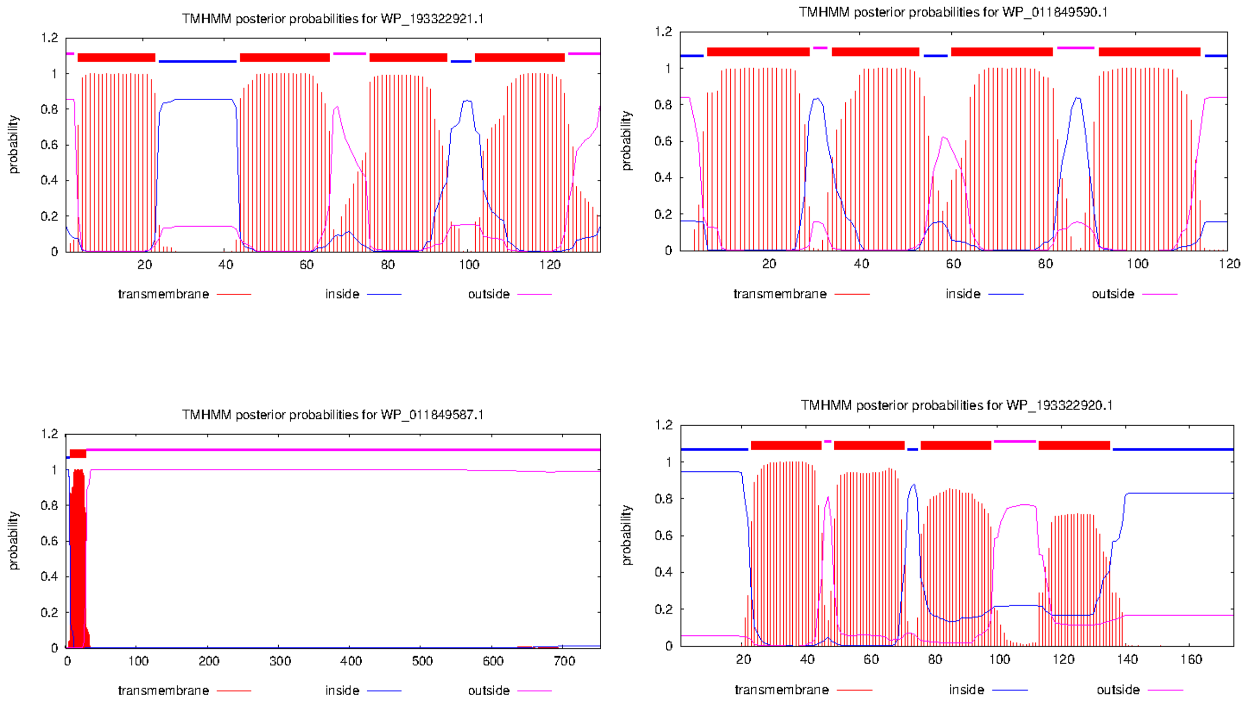


**Fig. S4.** Predicted transmembrane helices of Pcal_0908 (WP_193322921), Pcal_0907 (WP_011849590), Pcal_0904 (WP_011849587) and Pcal_0903 (WP_193322920). Predicted membrane spanning helical domains are indicated with red horizontal bars.

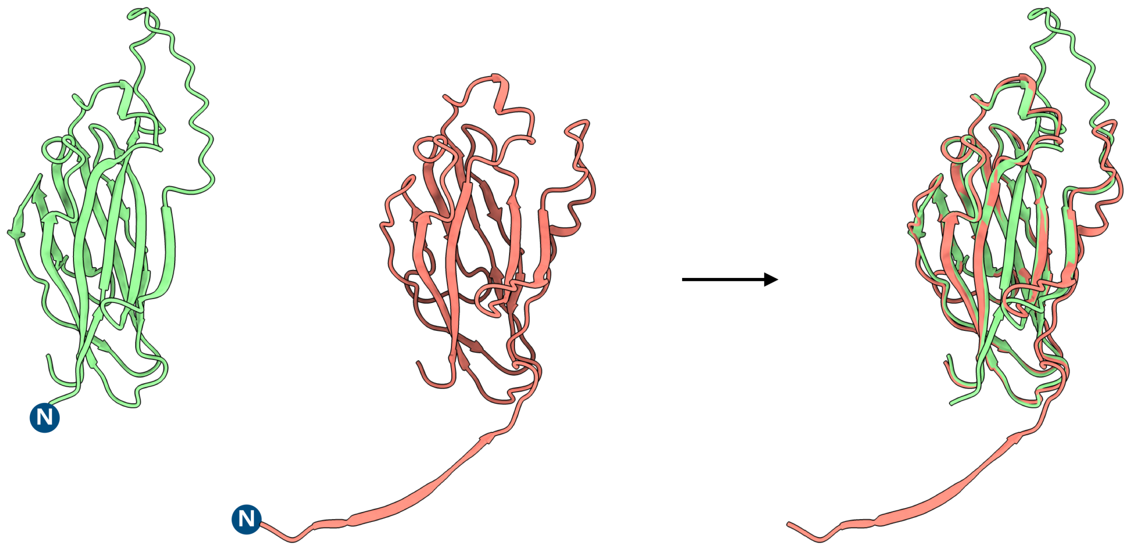


AlphaFold prediction | Cryo-EM structure | Alignment

**Fig. S5.** AlphaFold2 prediction vs cryo-EM determined ABP structure (single protein subunit)

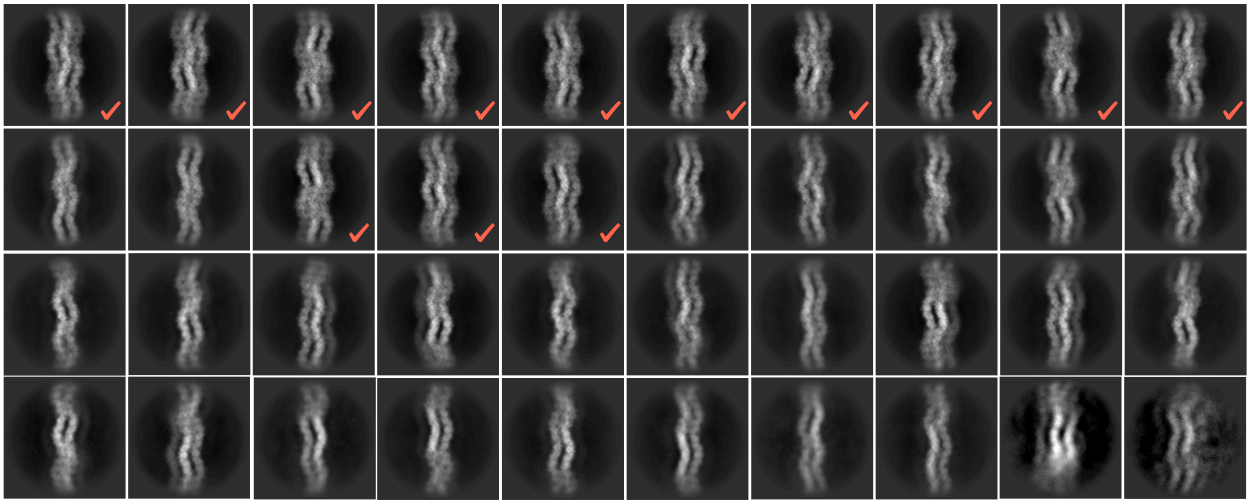

**Fig. S6.** 2D averages of ABP bundles. The red checkmarks indicate particles of the corresponding classes selected for 3D reconstruction.
